# PP1 promotes cyclin B destruction and the metaphase-anaphase transition by dephosphorylating CDC20

**DOI:** 10.1101/2020.06.22.164251

**Authors:** James Bancroft, James Holder, Zoë Geraghty, Tatiana Alfonso-Pérez, Daniel Murphy, Francis A. Barr, Ulrike Gruneberg

**Affiliations:** Sir William Dunn School of Pathology, University of Oxford, Oxford OX1 3RE; Department of Biochemistry, University of Oxford, Oxford OX1 3QU

## Abstract

Ubiquitin-dependent proteolysis of cyclin B and securin initiates sister chromatid segregation and anaphase. The anaphase promoting complex/cyclosome (APC/C) and its co-activator CDC20 form the main ubiquitin E3 ligase for these proteins. APC/C^CDC20^ is regulated by CDK1-cyclin B and counteracting PP1 and PP2A family phosphatases through modulation of both activating and inhibitory phosphorylations. Here we report that PP1 promotes cyclin B destruction at the onset of anaphase by removing specific inhibitory phosphorylation in the N-terminus of CDC20. Depletion or chemical inhibition of PP1 stabilises cyclin B and results in a pronounced delay at the metaphase-to-anaphase transition after chromosome alignment. This requirement for PP1 is lost in cells expressing CDK1-phosphorylation defective CDC20^6A^ mutants. These CDC20^6A^ cells show a normal spindle checkpoint response, but once all chromosomes have aligned rapidly degrade cyclin B and enter into anaphase in the absence of PP1 activity. PP1 therefore facilitates the metaphase-to-anaphase by promoting APC/C^CDC20^-dependent destruction of cyclin B in human cells.

## Introduction

Entry into and exit from mitosis is regulated by a conserved network of pathways controlling the activity and stability of the cyclin B dependent protein kinase (CDK1-cyclin B). CDK1-cyclin B activation is crucial for promoting entry into mitosis and the maintenance of the mitotic state (Nigg, 2001). Subsequently, destruction of cyclin B once all chromosomes have aligned is the key event permitting exit from mitosis (Clute and Pines, 1999). Cyclin B stability is controlled by a specific ubiquitin E3 ligase known as the anaphase promoting complex/cyclosome (APC/C) acting in concert with the ubiquitin-proteasome system (Alfieri et al., 2017; Sivakumar and Gorbsky, 2015; Watson et al., 2019). The APC/C initiates anaphase onset by ubiquitylating cyclin B and other substrates, tagging them for destruction by the proteasome (Irniger et al., 1995; King et al., 1995; Murray, 1995). CDK1-cyclin B modulates the activity of APC/C complexes towards different substrates through antagonistic phosphorylation of core APC/C subunits and two co-activators CDC20 and CDH1. CDK-phosphorylation of the core APC/C is necessary for CDC20-dependent ubiquitin ligase activity, whereas both CDC20 and CDH1 are inhibited by mitotic phosphorylation (Fujimitsu et al., 2016; Kraft et al., 2003; Kramer et al., 2000; Labit et al., 2012; Qiao et al., 2016; Yudkovsky et al., 2000; Zhang et al., 2016). Prior to mitosis when CDK1-cyclin B activity is low, the formation of active APC/C^CDC20^ complexes is unfavourable since the core APC/C subunits are not phosphorylated. Under these conditions an autoinhibitory segment within the APC1 subunit reduces binding of the CDC20 co-activator, and thus prevents ubiquitin-ligase activity towards cyclin B in late S-phase and G2 (Fujimitsu et al., 2016; Qiao et al., 2016; Zhang et al., 2016). CDH1 binding to the APC/C does not require activating phosphorylation of these core subunits (Zhang et al., 2016). CDH1 activation of the APC/C is prevented until late during exit from mitosis by two means. CDH1 is sequestered by an inhibitory factor EMI1 throughout S-phase and G2 (Hsu et al., 2002; Miller et al., 2006). Additionally, CDK1-dependent phosphorylation of CDH1 prevents it from binding to the core APC/C during mitosis (Kramer et al., 2000; Zachariae et al., 1998). Thus, CDK1 first promotes activation of APC/C^CDC20^ while simultaneously inhibiting APC/C^CDH1^. However, in the absence of further regulation, this arrangement would result in cyclin B destruction immediately upon entry into mitosis without a delay to allow for chromosome alignment and segregation, or cell division.

During mitosis, the core APC/C becomes phosphorylated and CDC20 would be expected to activate its ubiquitin ligase activity towards cyclin B immediately on mitotic entry. However, this is prevented by two further mechanisms. First, CDC20 is phosphorylated by CDK1 on entry into mitosis, reducing its affinity for the APC/C (Kramer et al., 2000; Yudkovsky et al., 2000). Second, CDC20 is sequestered into a diffusible inhibitor of the APC/C termed the mitotic checkpoint complex (MCC) until the process of chromosome alignment has been completed. MCC consists of four proteins, the checkpoint proteins MAD2, BUB3 and BUBR1, and the APC/C accessory subunit CDC20 (Fraschini et al., 2001; Hardwick et al., 2000; Lara-Gonzalez et al., 2012; Musacchio, 2015; Sudakin et al., 2001). Formation of the MCC is triggered at unattached or incorrectly attached kinetochores through the action of the spindle checkpoint kinase MPS1 (Abrieu et al., 2001). MPS1 promotes the accumulation of the MCC subunits at kinetochores through the phosphorylation of the outer kinetochore protein KNL1 (London et al., 2012; Shepperd et al., 2012; Yamagishi et al., 2012). MPS1-phosphorylated KNL1 acts as a landing platform for BUB1/BUB3 and BUBR1/BUB3 complexes (Overlack et al., 2015; Primorac et al., 2013). BUB1 is further phosphorylated by MPS1, initiating the recruitment of MAD1 (Ji et al., 2017; Qian et al., 2017; Zhang et al., 2017). Crucially, MPS1 catalyses the formation of MCC through phosphorylation of the C-terminus of the MAD1 spindle protein (Faesen et al., 2017; Ji et al., 2017). Further strengthening MCC production, phosphorylation by CDK1-cyclin B has been reported to bias CDC20 towards preferential incorporation into MCC rather than association with APC/C (D’Angiolella et al., 2003; Yudkovsky et al., 2000). Whether this is a direct effect of phosphorylation promoting incorporation of CDC20 into MCC, or an indirect effect of non-phosphorylated CDC20 preferentially binding to APC/C and thus changing the pool of CDC20 available for MCC formation, is unclear. Completing this regulatory circuit, CDK1-cyclin B1 directly aids the production of MCC by promoting the activation and recruitment of the checkpoint kinase, MPS1, to kinetochores (Alfonso-Perez et al., 2019; Hayward et al., 2019a; Hayward et al., 2019b; Morin et al., 2012; Vazquez-Novelle et al., 2014). This CDK1-cyclin B dependent cycle of MCC production and inhibition of APC/C thus prevents entry into anaphase until stable microtubule-kinetochore attachment has been achieved at all kinetochores.

CDK1-cyclin B therefore stabilises the mitotic state in two ways and in doing so creates a crucial requirement for regulated phosphatase activity in mitotic exit. In mitosis CDK1 inhibits APC/C activity through CDC20 and CDH1 phosphorylation and the spindle checkpoint pathway. Conversely, CDK1-cyclin B also promotes mitotic exit by phosphorylating and thereby activating APC/C (Kraft et al., 2003). Thus, CDC20 can only activate the APC/C in mitosis until the early stages of anaphase when APC/C phosphorylation is maintained. Maximal APC/C^CDC20^ activity in anaphase therefore requires differential dephosphorylation of the APC/C and CDC20. CDC20 must be dephosphorylated before the core APC/C is dephosphorylated, otherwise active APC/C^CDC20^ complexes would not form. Later dephosphorylation of CDH1 would then explain the formation and activity of APC/C^CDH1^. It has been suggested that the preference of the PP2A phosphatase for phosphorylated threonine compared to serine may explain differential dephosphorylation kinetics in mammalian cells. For CDC20 a number of the important phosphorylated regulatory sites are on threonine residues, whereas those on the APC/C and CDH1 are serine (Fujimitsu and Yamano, 2020; Hein et al., 2017). However, in other organisms there is good evidence that PP1 dephosphorylates CDC20 to active the APC/C at mitotic exit (Kim et al., 2017). In general, the modulation of CDK1-cyclin B mediated phosphorylation by counteracting phosphatases remains poorly understood, and this may reflect the involvement of multiple phosphatases acting on a diverse range of substrates. Both protein phosphatase 1 (PP1) and protein phosphatase 2A complexed to the B55 regulatory subunit (PP2A-B55) have been implicated as major CDK1-cyclin B opposing phosphatases (Cundell et al., 2016; Godfrey et al., 2017; McCloy et al., 2015; Schmitz et al., 2010; Wu et al., 2009). For PP2A-B55, unbiased proteomic screens have started identifying individual substrates as well as general motifs characterising PP2A-B55 targets in anaphase cells (Cundell et al., 2016; McCloy et al., 2015). Indeed, the aforementioned phosphorylation of MPS1 by CDK1-cyclin B1 is removed by PP2A-B55 during anaphase (Hayward et al., 2019a). In mammalian cells, PP2A-B55 has also been suggested as the enzyme responsible for CDC20 dephosphorylation (Hein et al., 2017). However, depletion of PP2A-B55 does not affect progression through the metaphase-to-anaphase transition (Cundell et al., 2013; Hayward et al., 2019a), which relies on normal APC/C^CDC20^ activation, making it unlikely that PP2A-B55 is the only phosphatase carrying out this job. For PP1, which has three isoforms in human cells, α, β and γ, unbiased substrate screens are still lacking, but substrates on chromatin and kinetochores have been identified (Cohen, 2002; Francisco and Chan, 1994; London et al., 2012; Nijenhuis et al., 2014; Nilsson, 2019; Qian et al., 2011; Wang et al., 2008; Yamashiro et al., 2008), including, in *C. elegans*, CDC20 (Kim et al., 2017).

Here we present evidence that PP1 is an important CDK1-counteracting CDC20 phosphatase in mammalian cells. By controlling APC/C^CDC20^-dependent destruction of cyclin B downstream of the spindle checkpoint, PP1 accelerates the metaphase-to-anaphase transition and contributes to the timing of mitotic exit.

## Results

### PP1 contributes to activation of APC/C during mitotic exit

Synchronous mitotic exit and entry into anaphase can be induced by chemical inhibition of the spindle checkpoint kinase MPS1 or the mitotic master kinase CDK1, permitting detailed biochemical analysis of mitotic exit (Cundell et al., 2013). Interestingly, inhibition of CDK1 results in accelerated destruction of cyclin B1 when compared to inhibition of the checkpoint kinase MPS1 (Figure S1A and S1B) (Cundell et al., 2013). This agrees with the notion that CDK1-cyclin B directly opposes APC/C activation independently of the MPS1-dependent spindle checkpoint pathway. In principle, this could be explained by CDK1-counteracting phosphatases of the PP1 and PP2A families. However, previous work showed that silencing of PP2A-B55 does not delay the metaphase-to-anaphase transition in the absence of chromosome segregation errors (Cundell et al., 2013; Hayward et al., 2019a). By contrast, PP2A-B56 is active throughout mitosis and is essential for chromosome alignment and checkpoint signalling making it difficult to probe any downstream role at the metaphase-to-anaphase transition (Espert et al., 2014; Nilsson, 2019). We therefore focussed on the role of PP1, in part because CDK1 inhibition results in accelerated dephosphorylation of the inhibitory T320 residue in PP1 with kinetics that parallel those for cyclin B destruction (Figure S1A and S1B). PP1 is phosphorylated and inhibited by CDK1 on a conserved C-terminal threonine, T320 in the PP1α isoform and the equivalent residues in the PP1β and PP1γ isoforms (Dohadwala et al., 1994; Goldberg et al., 1995; Kwon et al., 1997). The dephosphorylation of this residue is thought to be an auto-dephosphorylation event and can therefore be used as surrogate measure of PP1 activity (Wu et al., 2009).

To test whether PP1 activity contributes to the accelerated destruction of cyclin B1 we performed a biochemical analysis of mitotic exit in the presence of the highly specific small molecule PP1 inhibitor tautomycetin which inhibits all three PP1 isoforms but not PP2A (Choy et al., 2017; Hayward et al., 2019c). Tautomycetin prevented dephosphorylation of PPP1CA-pT320 (Figure 1A and 1B), and the half-life of pT320 increased from ∼2 min in the control to >60 min (Figure 1C). Consistent with the idea that PP1 plays a role in the regulation of cyclin B stability in mitotic exit, cyclin B1 destruction was delayed in the presence of tautomycetin (Figure 1A and 1B), and the half-life increased from 4 min to 12 min (Figure 1D).

**Figure 1.**
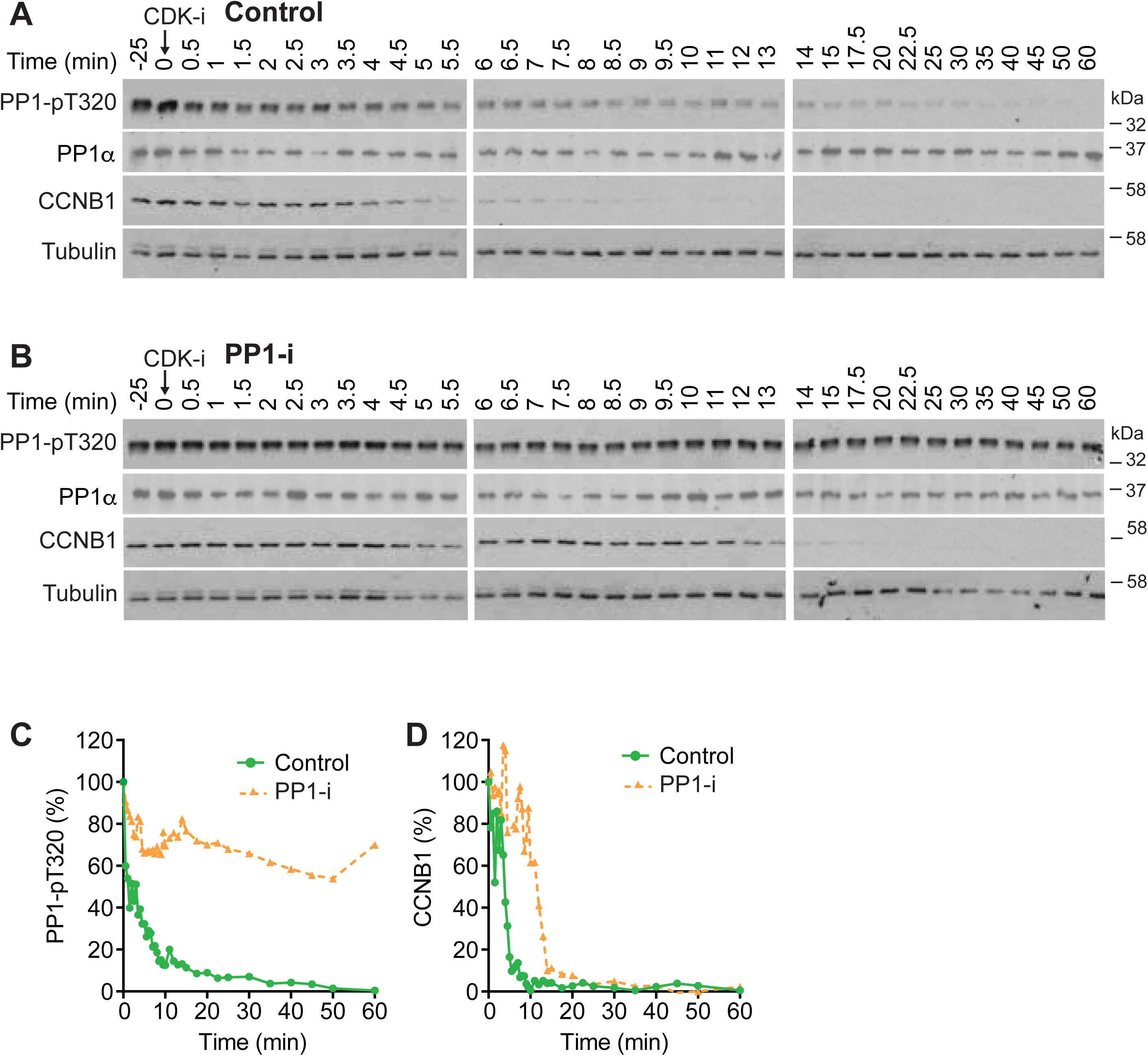
PP1 counteracts CDK1-regulation of APC/C during mitotic exit. **(A)** HeLa cells were arrested in mitosis and incubated for 25 min with either DMSO (Control) or **(B)** 5 µM tautomycetin PP1 inhibitor (PP1-i). CDK1 was then inhibited (CDK-i) using 5 µM flavopiridol and samples of the culture taken at the times indicated and Western blotted for the indicated markers. **(C)** Densitometric quantification of PP1α-pT320 and **(D)** CCNB1 is plotted in the line graphs.

To identify which specific isoform of PP1 was responsible for this effect, HeLa cells were depleted of PP1 catalytic subunits, arrested in mitosis and then treated with CDK1 inhibitors. Since PP1α (PPP1CA) and PP1γ (PPP1CC) are reported to have overlapping functions during mitosis (Liu et al., 2010; Trinkle-Mulcahy et al., 2006), these two catalytic subunits were depleted simultaneously (siPP1α/γ). PP1β (PPP1CB) which has been reported to have a role distinct from PP1α and PP1γ (Kiss et al., 2019; Matsumura et al., 2011; Yamashiro et al., 2008), was knocked down separately. High-resolution time courses with cell samples every 30 sec were then collected in the window of cyclin B destruction. In this assay, cyclin B1 and securin were rapidly degraded after CDK1 inhibition with a t_1/2_ of 11 mins (Figure 2A and 2E), whereas co-depletion of PP1α and PP1γ slowed the rate of cyclin B1 destruction to a t_1/2_ of 17 mins (Figure 2B and 2E). PP1β depletion did not alter the kinetics of cyclin B or securin destruction when compared to the control (Figure 2C and 2E). Efficient PP1 depletion was confirmed for the different conditions by western blotting (Figure 2D). We conclude that PP1 activity reverses CDK1-dependent phosphorylation of proteins normally limiting APC/C^CDC20^ activity towards cyclin B1 and securin at the metaphase-to-anaphase transition (Figure 2F).

**Figure 2.**
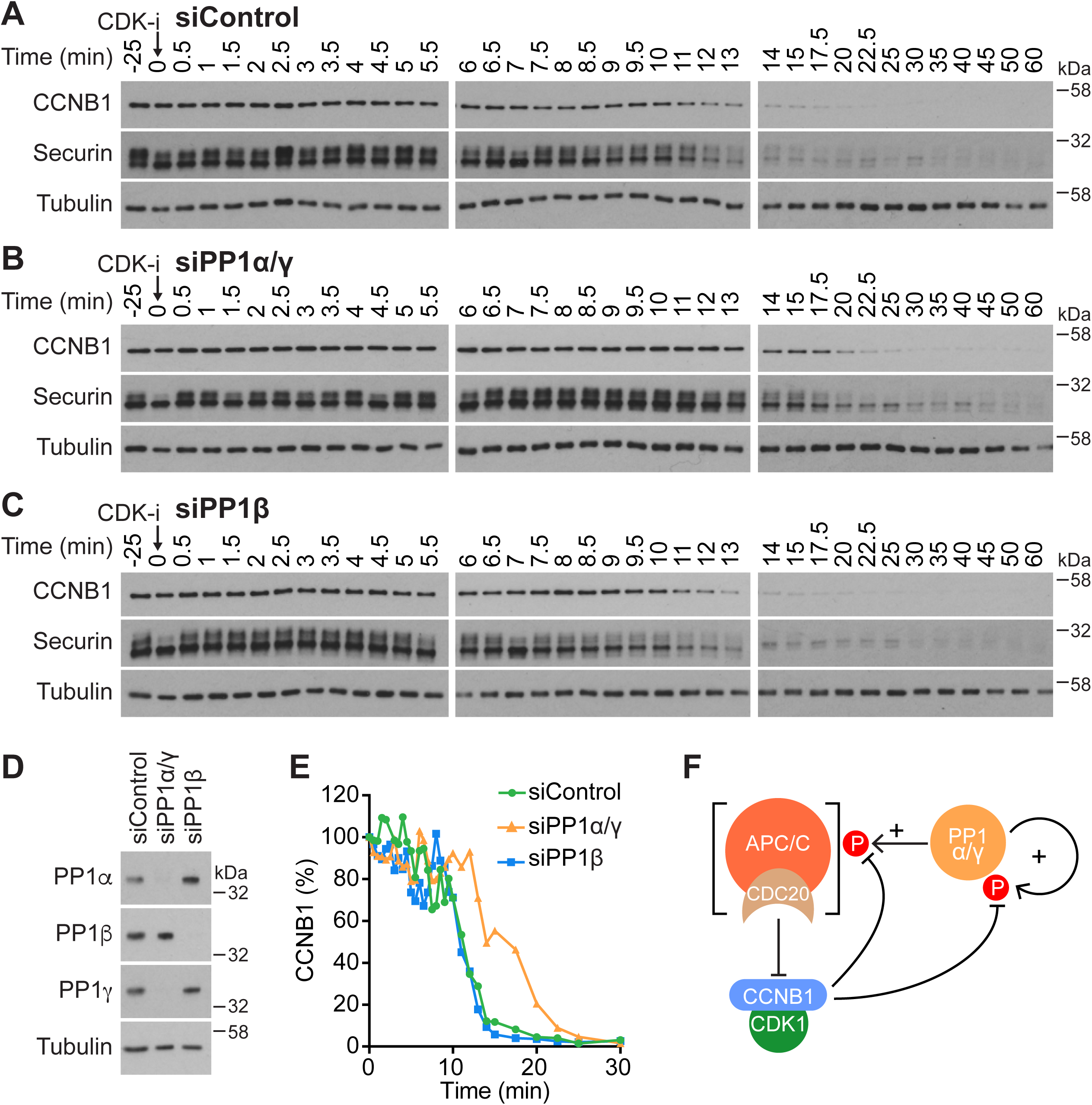
Identification of the major PP1 isoforms counteracting CDK1-regulation of APC/C. **(A)** HeLa cells were treated with either control siRNA or depleted for catalytic subunits of **(B)** PP1α/γ or **(C)** PP1β, and then arrested in mitosis. Mitotic exit was then triggered by CDK1 inhibition (CDK-i), and samples taken for Western blot at the times indicated in the figure. **(D)** Western blot confirming efficient depletion of PP1 catalytic subunits. **(E)** Densitometric quantification of CCNB1 levels across the three conditions. **(F)** A schematic depicting the potential role of PP1 in APC/C activation.

### PP1 activity is required to trigger prompt cyclin B1 degradation

To investigate the role of PP1 at the metaphase-to-anaphase transition without the need to use CDK1-inhibition, we performed a single cell analysis using HeLa cells expressing GFP-tagged cyclin B1 (CCNB1) from the endogenous promoter (Alfonso-Perez et al., 2019). This allowed us to follow cyclin B1 degradation in individual control cells passing through mitosis and into anaphase, and to compare control with PP1α/γ- or PP1β-depleted cells. In agreement with the biochemical data showing PP1 is required for rapid cyclin B destruction, cells depleted for PP1α/γ showed delayed passage through mitosis. PP1α/γ depleted cells took >100 min to reach anaphase from nuclear envelope breakdown compared to 60-70 min for control or PP1β depleted cells (Figure 3A). This could be largely attributed to a delay in the time taken to proceed from the completion of the metaphase plate to anaphase (Figure 3B). In good agreement with previous observations (Clute and Pines, 1999), cyclin B1 was quickly degraded once chromosomes had aligned and established a metaphase plate in control cells, and this was followed by segregation of the chromosomes (Figure 3C and 3D). However, depletion of PP1α and PP1γ delayed cyclin B1 destruction (Figure 3C and 3E), resulting in a significantly increased overall length of mitosis and specifically an extended metaphase-to-anaphase transition (Figure 3A and 3B). A sub-population of cells displayed wild type kinetics for cyclin B1 destruction, possibly due to incomplete or variable levels of PP1α/γ depletion. Depletion of PP1β had no obvious effect on cyclin B destruction compared to the control condition (Figure 3B and 3F).

**Figure 3.**
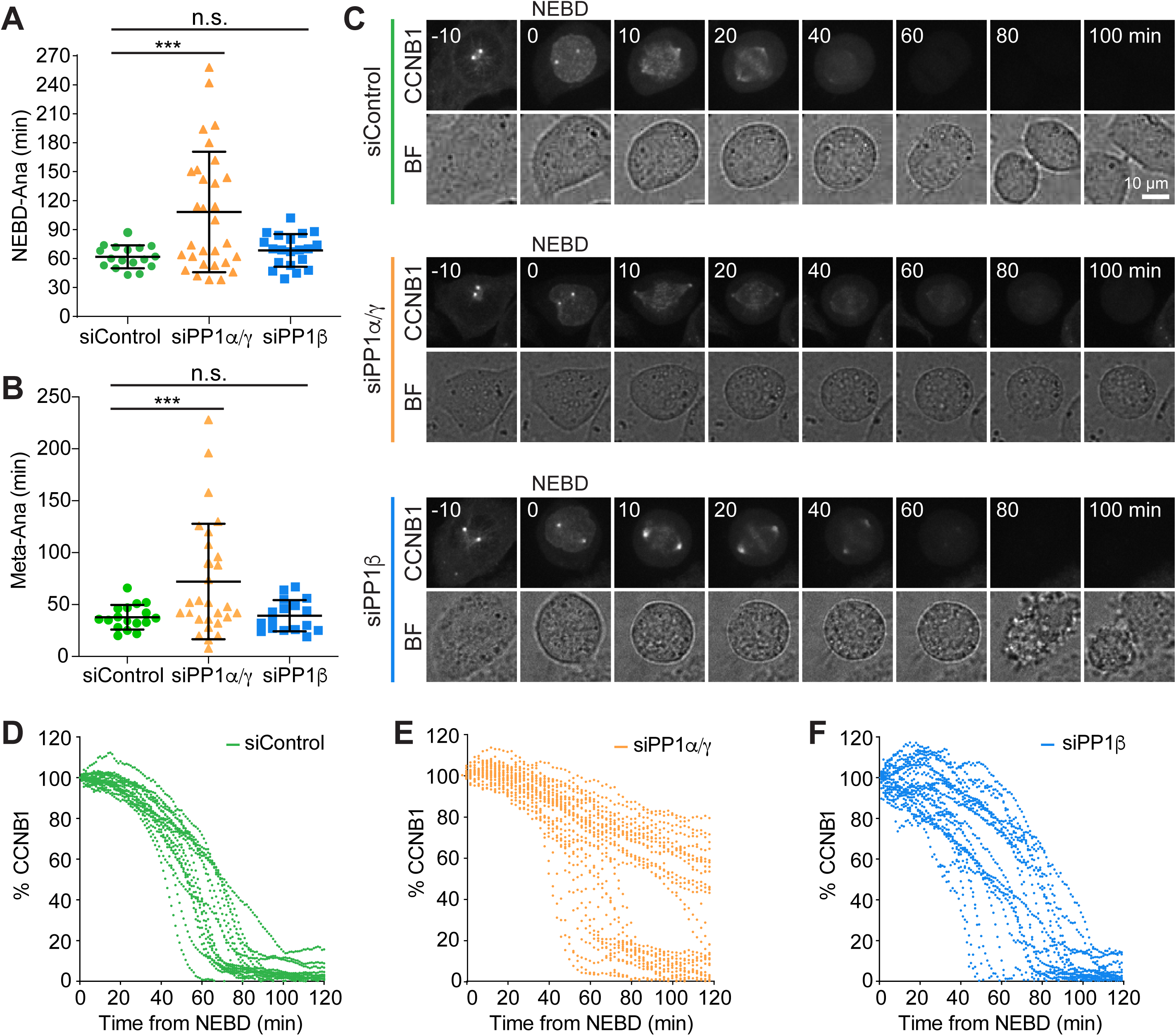
PP1 is needed for rapid destruction of cyclin B at the metaphase-anaphase transition. **(A)** Comparison of the time spent in mitosis or **(B)** to enter anaphase following completion of a metaphase plate in siControl, siPP1α/γ or siPP1β depletion for CRISPR-tagged CCNB1-GFP HeLa cells. Scatter plots of mean ± SD are shown (siControl n=17, siPP1α/γ n=30, siPP1β n=20). **(C)** Live cell imaging of CCNB1-GFP with times shown in minutes. Percentage of CCNB1-GFP fluorescence following NEBD in **(D)** siControl, **(E)** siPP1α/γ and **(F)** siPP1β cells. Images shown are maximum intensity projections, quantification was carried out on sum intensity projections and the single cell traces plotted as a function of time from NEBD in the graphs.

PP1 therefore promotes cyclin B destruction at the metaphase-to-anaphase transition in unperturbed mitosis, confirming the data obtained under CDK1-inhibited conditions.

### PP1 activity promotes timely progress from metaphase to anaphase

PP1 together with PP2A-B56 has been reported to counteract the MPS1-dependent phosphorylation of MELT motifs in KNL1 (London et al., 2012; Meadows et al., 2011; Nijenhuis et al., 2014; Rosenberg et al., 2011). Removal of PP1 would therefore be expected to delay silencing of the spindle assembly checkpoint. This could explain, either fully or in part, the delayed degradation of cyclin B1 upon PP1 inhibition and in PP1α/γ depleted cells. Alternatively, in *C. elegans* it has been found that PP1 can also play a role in APC/C activation downstream of the spindle checkpoint (Kim et al., 2017). To investigate both these possibilities and define when PP1 is acting, we imaged HeLa cells stably co-expressing CCNB1-mCherry from its endogenous promoter and a GFP-tagged version of the spindle checkpoint protein MAD2. We focussed on asking if the period from spindle checkpoint silencing to the onset of anaphase was increased.

The defining hallmark of spindle checkpoint silencing is the loss of all MAD2-positive kinetochores. In control cells, complete chromosome alignment coincided with disappearance of the last MAD2-positive kinetochore, rapidly followed by loss of the CCNB1-mCherry signal and chromosome segregation (Figure 4A and 4D). Inhibition of PP1 with tautomycetin delayed cyclin B destruction following loss of the last MAD2-positive kinetochore (Figure 4B and 4D). The effect observed resembled a CDC20 depletion, in which spindle checkpoint silencing proceeded normally but cyclin B degradation was not initiated (Figure 4C and 4D, Figure S3D). At 50 minutes after checkpoint silencing, as judged by the disappearance of GFP-MAD2, PP1 inhibited cells retained the same level of cyclin B as CDC20 depleted cells which are unable to target cyclin B for destruction (Figure 4E).

**Figure 4.**
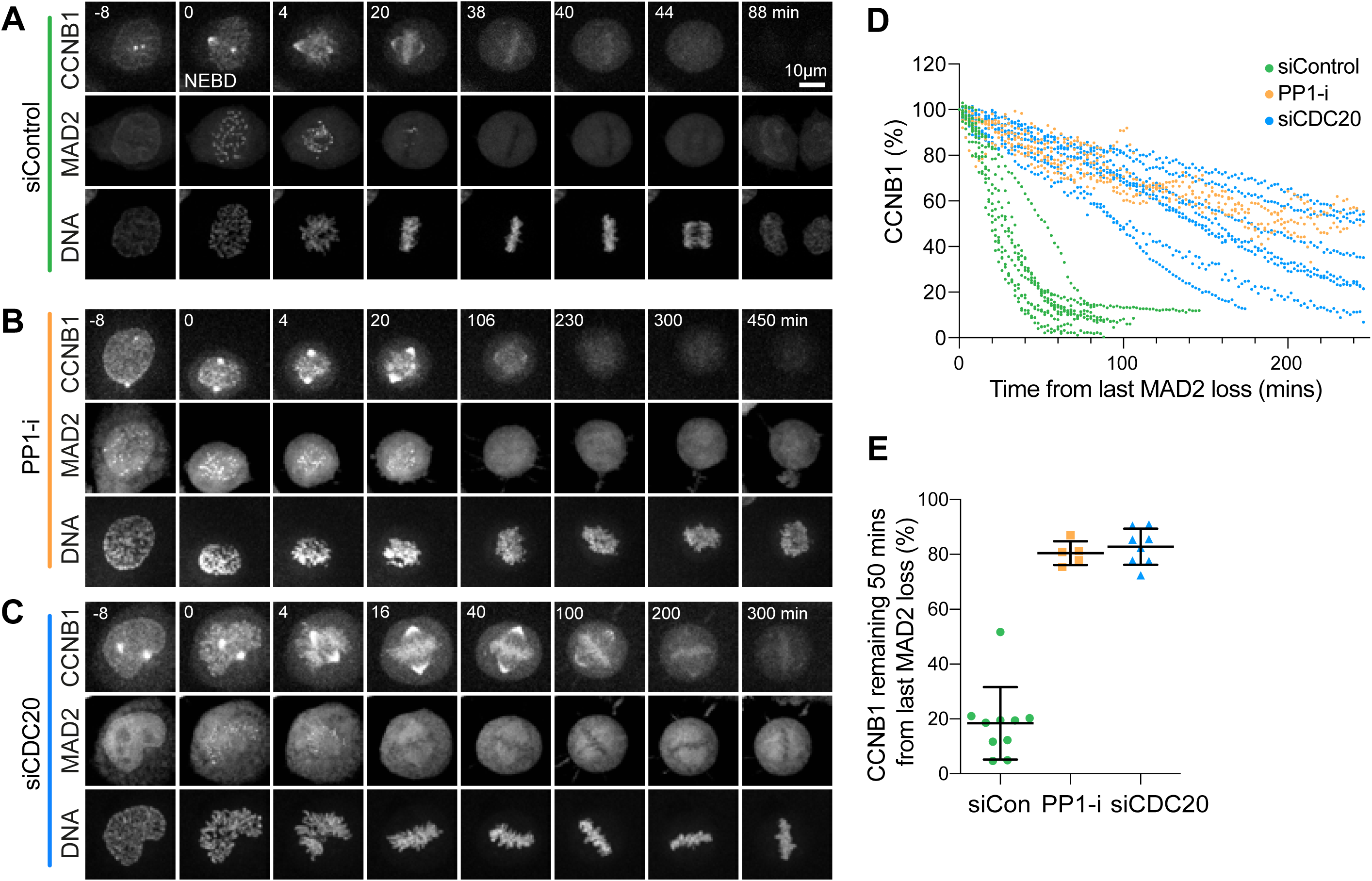
PP1 inhibition prevents rapid cyclin B destruction downstream of the spindle checkpoint. **(A)** HeLa CCNB1-mCherry GFP-MAD2 cells treated with control siRNA, **(B)** 5 µM PP1 inhibitor tautomycetin (PP1-i) for 30 min prior to the start of imaging, or **(C)** siCDC20 were imaged for 12 hours at 2 min intervals. Chromosome congression was monitored using SiR-Hoechst, and SAC silencing was inferred from the loss of GFP-MAD2 from the kinetochore. Degradation of endogenously tagged CCNB1 indicated APC/C activation. Representative time course images are shown. **(D)** CCNB1 levels were measured as a function of time from the point at which the last MAD2-positive kinetochore was observed for siControl, PP1-i and siCDC20 cells. CCNB1 levels were set to 100% at that point and plotted for single cells in the graph. **(E)** Scatter plots for the percentage of CCNB1 signal remaining at 50 mins after observation of the last MAD2-positive kinetochore under each experimental condition in (A-C) show the mean ± SD (siControl, n=10; PP-i (5 µM tautomycetin) n=5; siCDC20, n=8).

We then examined the relationship between checkpoint silencing and cyclin B destruction in PP1α/γ depleted cells, focussing first on checkpoint silencing. Efficient depletion of PP1 catalytic subunits was first confirmed by Western blotting and immunofluorescence analysis of cells expressing GFP-tagged PP1 catalytic subunits (Figure S1E-G). This analysis revealed that MAD2 positive kinetochores were observed for a longer period in PP1-inhibited or PP1α/γ -depleted cells than in control cells (Figure S2A-S2C). However, PP1-inhibited or PP1α/γ -depleted cells were able to silence the checkpoint and enter anaphase with a delay (Figure S2B and S2C). In agreement with the idea that checkpoint silencing and cyclin B destruction are independently delayed, both the times taken for spindle checkpoint silencing to complete and the time from loss of the last MAD2 signal to the onset of anaphase were increased (Figure S2F). By contrast, PP2A-B56 depleted cells arrested with sustained MAD2-positive kinetochores and failed to enter anaphase (Figure S2D and S2E). These observations are consistent with a major role for PP2A-B56 in the spindle checkpoint and error correction pathways, with a contribution by PP1 to the complete silencing of the spindle checkpoint (Espert et al., 2014; Nijenhuis et al., 2014).

We then measured the kinetics of cyclin B1 destruction in PP1 depleted cells to test if this was delayed even after the spindle checkpoint had been silenced. In comparison to control cells, cyclin B1 destruction was delayed after the loss of MAD2-positive kinetochores in PP1α/γ depleted cells (Figure 5A, 5B and 5D). At 50 mins after the loss of the last MAD2-positive kinetochore, control cells had completed destruction of cyclin B1 (Figure 5D and 5E). By contrast, PP1-depleted cells retained ∼80% of the maximal level of cyclin B1 measured at nuclear envelope breakdown (NEBD) (Figure 5D and 5E). Importantly, PP2A-B55 depletion did not change the kinetics of the metaphase-to-anaphase transition or cyclin B1 degradation (Figure 5C-5E, Figure S3D).

**Figure 5.**
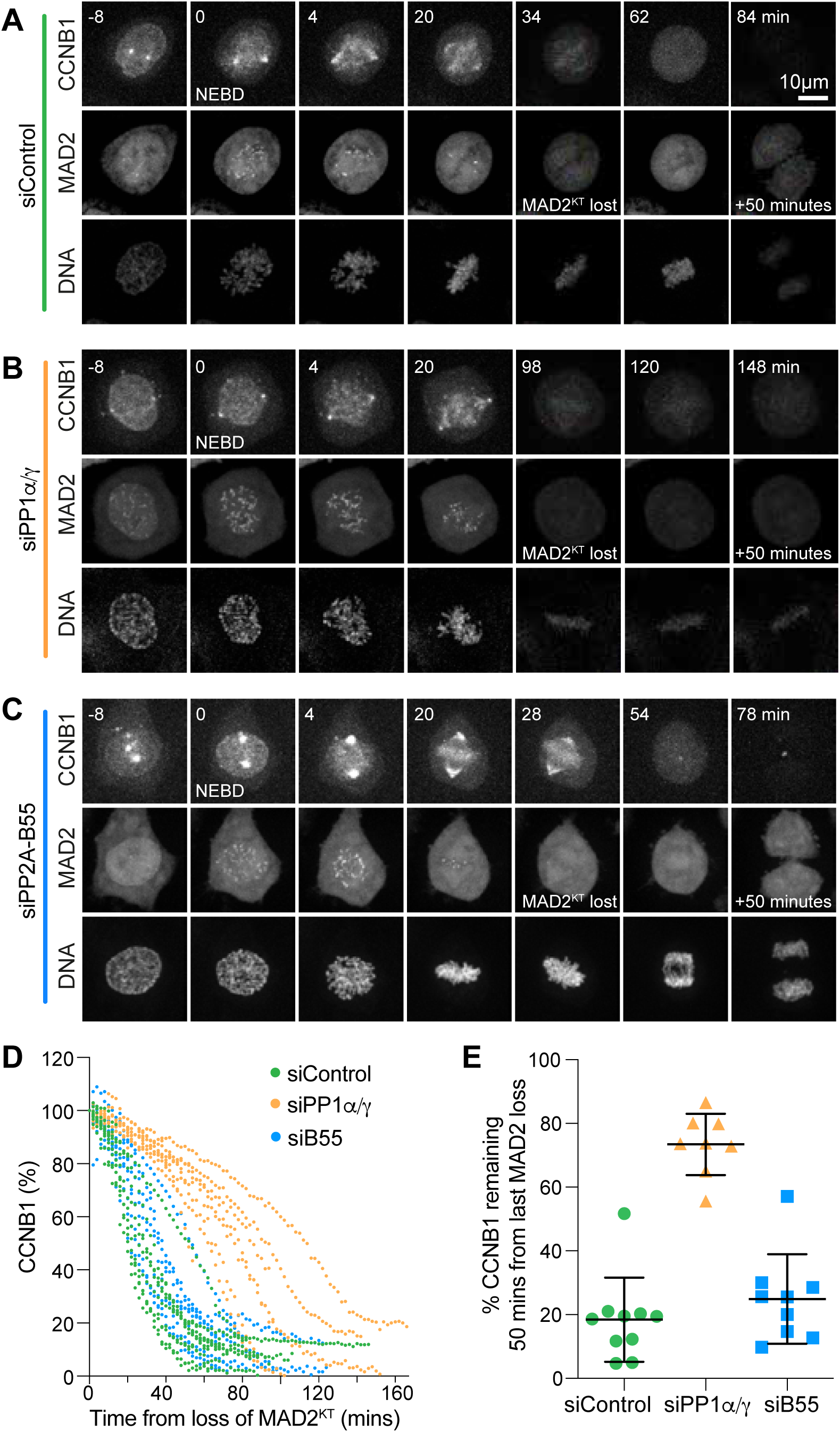
PP1α/γ are required for rapid cyclin B destruction downstream of the spindle checkpoint. **(A)** HeLa CCNB1-mCherry GFP-Mad2 cells, treated with control, **(B)** siPP1α/γ or **(C)** siPP2A-B55 for 60 hours were imaged for 12 hours at 2 min intervals. DNA was visualised with SiR-DNA. **(D)** Quantitation of the CCNB1 levels of individual cells shown in (A). **(E)** The CCNB1 signal remaining at 50 min in siControl, siPP1α/γ or siPP2A-B55 was plotted as mean ± SD (siControl n=10, siPP1α/γ n=8, siPP2A-B55 n=9).

To further confirm that the delay in cyclin B degradation observed in cells depleted of PP1 activity was independent of spindle checkpoint activity, we artificially silenced spindle checkpoint signalling by addition of the specific MPS1 inhibitor AZ3146 in cells approaching metaphase, and then measured the decline of cyclin B levels in control and PP1 depleted cells (Figure 6A and 6B). In all control and PP1 depleted cells, MPS1 inhibitor addition resulted in a rapid loss of kinetochore associated MAD2 within 2-4 min (Figure 6A). Cyclin B degradation, however, was strongly delayed in PP1 depleted cells in comparison to control cells, suggesting that this was not dependent on MPS1 mediated spindle checkpoint activity.

**Figure 6.**
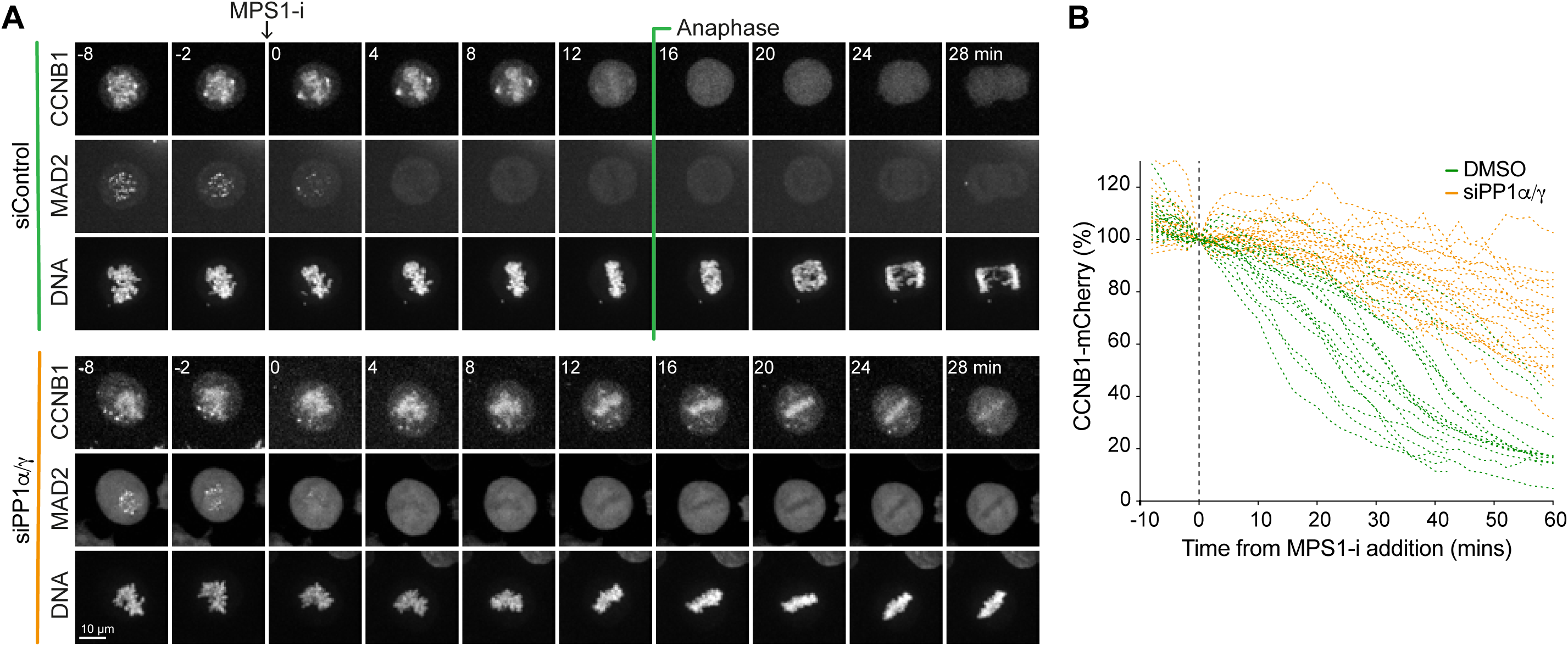
In the absence of PP1α/γ, cyclin B degradation is delayed when the spindle checkpoint is inactivated by MPS1 inhibition. **(A)** HeLa CCNB1-mCherry GFP-MAD2 cells treated with control or siPP1α/γ, were imaged progressing through mitosis and treated with MPS1 inhibitor AZ3146 (MPS1-i) when they approached metaphase. Loss of GFP-MAD2 and kinetics of CCNB1-mCherry degradation were monitored. **(B)** CCNB1-mCherry levels in single cells are plotted for siControl and siPP1α/γ cells undergoing mitotic exit. The time of MPS1-i addition (t=0) is marked with a dashed line.

Taken together, these results confirm that PP1 plays an important role in mediating the metaphase-to-anaphase transition, by contributing to spindle checkpoint silencing and through subsequent downstream regulation of cyclin B destruction.

### PP1 dephosphorylates the N-terminus of CDC20

The results presented so far support the view that PP1 activity is required to trigger the rapid APC/C-dependent destruction of cyclin B1 downstream of the spindle checkpoint. The most parsimonious explanation for this observation is that one or more of the core components of the APC/C or an APC/C co-activator are dephosphorylated by PP1. Because CDC20 has been described as a PP1 target at the metaphase-to-anaphase transition in *C. elegans* (Kim et al., 2017), we decided to test the hypothesis that CDC20 may also be a critical PP1 target in human cells. CDC20 is phosphorylated by CDK1 on six amino acids in the N-terminus of the protein (Figure 7A) (Labit et al., 2012; Yudkovsky et al., 2000). To investigate the kinetics of CDC20 dephosphorylation at the metaphase-to-anaphase transition in the presence and absence of PP1 activity, we first used PhosTag SDS-PAGE to visualise the mitotic phosphorylation of CDC20. PhosTag binds to phosphate groups on proteins and results in enhanced separation of the phosphorylated species on SDS-PAGE (Kinoshita et al., 2006). Comparison of control and PP1-inhibited cells lysates, as well as control and PP1 depleted cell lysates on PhosTag gels, revealed a pronounced PP1-dependent downshift in CDC20 as cells progressed from mitosis into anaphase following MPS1 inhibition (Figure 7B and 7D-7F). Western blotting with a CDC20 antibody specific for phosphorylated T70 confirmed that this effect was due to retention of phosphorylation (Figure 7C; Figure S3A-S3C). To allow a direct comparison to the biochemical data shown in Figure 1, cells were also forced into anaphase by CDK1 inhibitor treatment. Under these conditions CDC20 T70 was rapidly dephosphorylated in control cells, but not in the PP1 inhibitor treated cells (Figure 7G). Together, these results suggest that PP1 is an important CDC20 phosphatase at the metaphase-to-anaphase transition and identify T70 as a key diagnostic site for this regulation.

**Figure 7.**
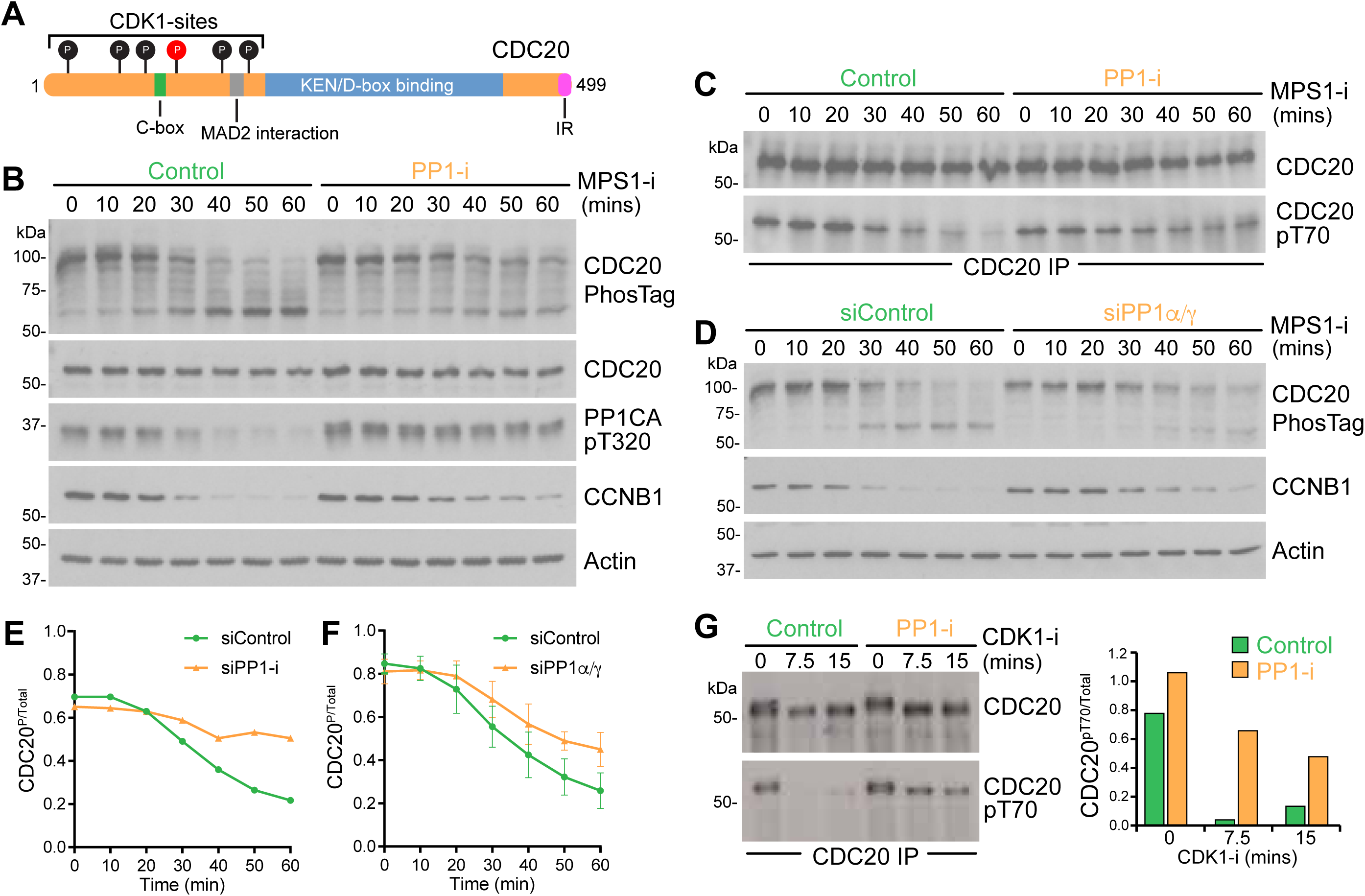
PP1 dephosphorylates the N-terminal region of CDC20. **(A)** Schematic representation of CDC20 structure. Six N-terminal CDK1 sites are indicated, the pT70 site is marked with in red. **(B)** Synchronous progression of mitotic HeLa cells pre-treated with DMSO or PP1 inhibitor into anaphase was triggered with MPS1 inhibitor and samples were collected every 10 min. Cell cycle progression was monitored by cyclin B1 Western blot; CDC20 phosphorylation state by PhosTag gel; and PP1 activation status by Western blot for PP1-pT320. Actin was used as a loading control. **(C)** CDC20 was immunoprecipitated from cells synchronized as in (B). The immunoprecipitates were blotted for pT70 modified CDC20 and the total amount of CDC20. **(D)** Synchronous progression of control depleted or PP1α/γ depleted mitotic HeLa cells into anaphase was triggered with MPS1 inhibitor and samples were collected as in (B). **(E)** Densitometric quantification of phospho-CDC20 (top band in the PhosTag blot) relative to total CDC20 in (B) and **(F)** in (D) is plotted in the line graphs. **(G)** Synchronous progression of mitotic HeLa cells pre-treated with DMSO or PP1 inhibitor into anaphase was triggered with CDK1 inhibitor and samples were collected every 7.5 min. CDC20 was immunoprecipitated and the immunoprecipitates blotted for pT70. The CDC20-pT70 levels relative to total CDC20 were blotted as bar graphs.

### Phosphorylation defective CDC20 obviates the need for PP1

If CDC20 is a major PP1 target at the point of anaphase onset, then a CDK1-phosphorylation resistant form of CDC20 should be able to rescue the cell cycle delay observed upon PP1 depletion or inhibition. To test this idea, endogenous CDC20 was depleted and replaced with either GFP-CDC20^WT^ or with GFP-CDC20^6A^, lacking the N-terminal CDK1-phosphorylation sites. In the presence of PP1, both versions of CDC20 resulted in a rescue of the cell cycle arrest observed upon CDC20 depletion (Figure 8A and 8B). Interestingly, cells expressing GFP-CDC20^6A^ passed through mitosis slightly faster than cells expressing CDC20^WT^, the half-life of mitosis from NEBD was shortened from 80 to 60 minutes, similar to results obtained with a phosphorylation deficient CDC20 mutant in *C. elegans* (Kim et al., 2017) (Figure 8E). These effects were not due to differences in CDC20 levels, since these were comparable for all conditions (Figure 8F). This dominant effect on APC/C activity in the presence of endogenous wild-type CDC20 is consistent with the idea that inhibitory regulation is attenuated or lost for the GFP-CDC20^6A^ mutant. When PP1 was depleted, a cell cycle delay at the metaphase-to-anaphase transition was observed in cells rescued with CDC20^WT^ (Figure 8C and 8D). This delay was alleviated by replacement of the endogenous CDC20 with GFP-CDC20^6A^ (Figure 8C and 8D). However, the half-life of mitosis was still extended compared to cells expressing normal levels of PP1 and wild-type CDC20 (Figure 8E). This agrees with the notion that PP1 has other targets affecting the onset of anaphase, for example in the spindle checkpoint pathway. Because of this it was important to test if CDC20^6A^ was defective for spindle checkpoint function. When GFP-CDC20^6A^ cells were treated with both high and low doses of nocodazole, the cells showed a normal cell cycle arrest (Figure 8G). This shows that the ability to trigger a spindle checkpoint response, that is MCC formation and inhibition of APC/C^CDC20^, was not grossly compromised by expression of phosphorylation resistant GFP-CDC20^6A^. We therefore conclude that CDC20 is one of the key targets for dephosphorylation by PP1 at the metaphase-to-anaphase transition, independent of its role in the spindle checkpoint. The rescue of both rapid cyclin B destruction and cell cycle progression in PP1 depleted cells by CDC20^6A^ is thus a reflection of the need for PP1-mediated dephosphorylation of CDC20 to enable timely progression into anaphase.

**Figure 8.**
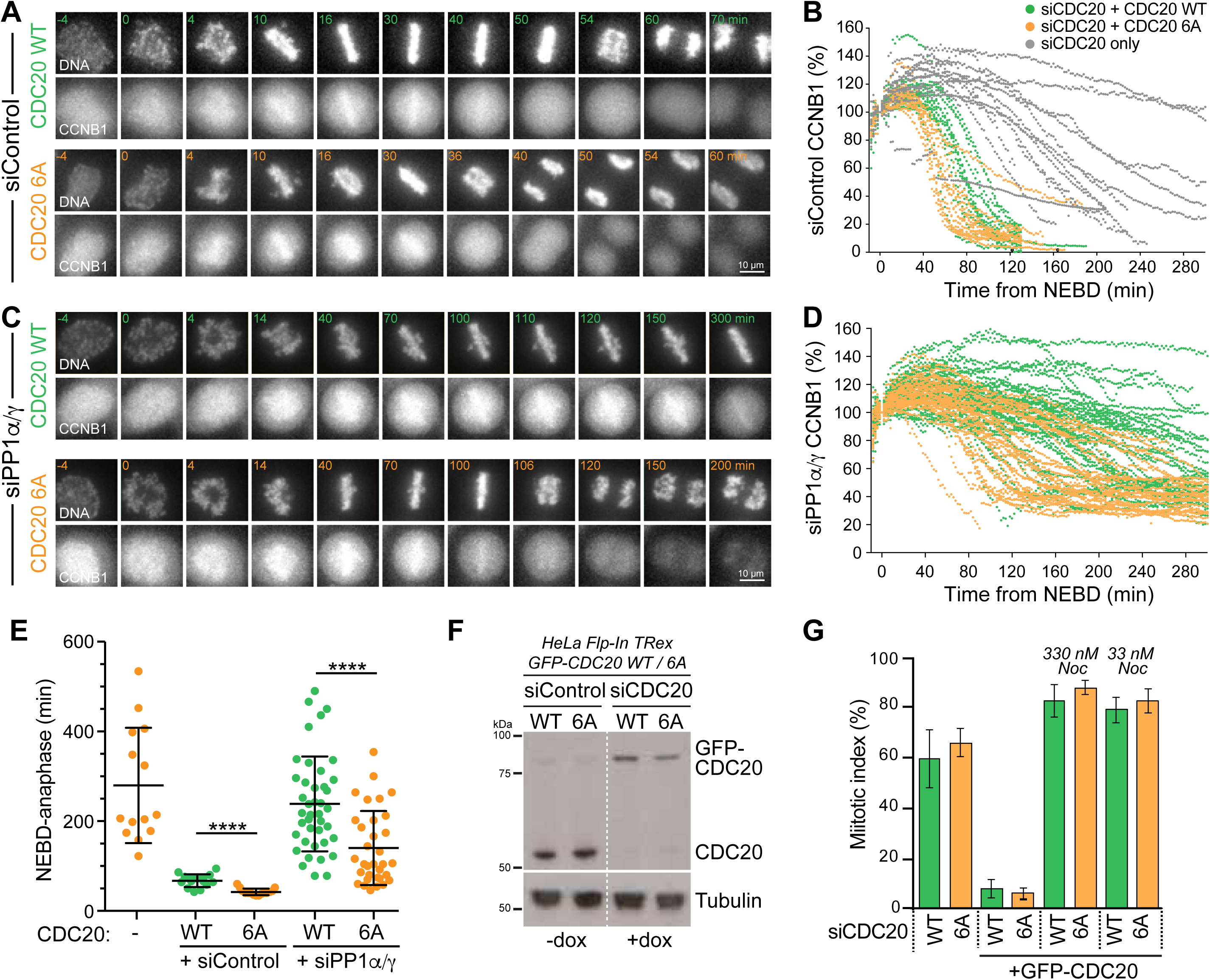
Phosphorylation resistant CDC20^6A^ bypasses the requirement for PP1 in rapid cyclin B destruction on mitotic exit. **(A and B)** HeLa Flp-In TRex cells expressing CCNB1-mCherry were depleted of endogenous CDC20 and induced to express either GFP-CDC20^WT^ or GFP-CDC20^6A^, in the presence or **(C and D)** absence of PP1α/γ. Cells cycle progression and CCNB1 levels were then followed by live cell imaging. DNA was visualised with SiR-DNA. CCNB1 levels in individual cells are plotted in the line graphs in (B) and (D). **(E)** Scatter plots showing the mean time ± SD at which the cells imaged entered anaphase or the end of the movie was reached (siControl: n= 20 for GFP-CDC20^WT^ cells (green), n=15 for GFP-CDC20^6A^ (orange), n=12 for uninduced cells (grey). siPP1α/γ n=41 for GFP-CDC20^WT^, n= 33 for GFP-CDC20^6A^). **(F)** Western blot of cells depleted of endogenous CDC20 and expressing GFP-CDC20 as in (A), arrested with 330 nM nocodazole for 14 hours. **(G)** Cells depleted for endogenous CDC20 and expressing GFP-CDC20^WT^ or 6A were treated with nocodazole at the indicated concentrations for 14 hours, and the mitotic index was plotted.

## Discussion

### CDC20 regulation by PP1

The metaphase-to-anaphase transition is a perilous stage during eukaryotic mitosis, marking the point of no return for correction of errors in chromosome alignment (Nasmyth, 2001). Mechanistically this is clearly understood since the release of APC/C inhibition at this point results in the activation of separase and consequently the loss of sister chromatid cohesion at all chromosomes simultaneously (Uhlmann, 2001). To ensure the synchronised separation and segregation of all the chromosomes, it is therefore crucial that the APC/C can be rapidly activated, yet is not triggered prematurely. Untimely APC/C activity at the metaphase-anaphase transition is prevented by both the spindle assembly checkpoint and phosphorylation of the APC/C coactivator CDC20 (Fujimitsu et al., 2016; Hein et al., 2017; Hein and Nilsson, 2016; Labit et al., 2012; Zhang et al., 2016). Phosphorylation of the N-terminal region of CDC20 has two notable consequences. First, to reduce the affinity of CDC20 for APC/C, and second to bias its incorporation into MCC rather than APC/C (D’Angiolella et al., 2003; Hein and Nilsson, 2016). Thus, CDC20 has to be dephosphorylated in order to achieve full APC/C activity in exit from mitosis (Labit et al., 2012). Based on the data presented here, we conclude that in human cells PP1 is needed for timely CDC20 dephosphorylation at the metaphase-to-anaphase transition. Supporting the view that this is a conserved mechanism of APC/C regulation, PP1 was shown to play a similar role in *C. elegans* (Kim et al., 2017).

### Local pools of PP1 in checkpoint signalling and APC/C regulation

A number of distinct pools of PP1 have been reported to contribute to the regulation of the metaphase-to-anaphase transition in mammalian cells. PP1 catalytic subunits bind to regulatory subunits via conserved short, linear motifs, the best characterised of which is the RVxF motif (Egloff et al., 1997; Terrak et al., 2004). Several RVxF-dependent PP1-interaction partners have been identified at attached kinetochores, including the outer kinetochore proteins KNL1 and Astrin and the mitotic motor proteins CENP-E and KIF18A (Conti et al., 2019; De Wever et al., 2014; Hafner et al., 2014; Kim et al., 2010; Liu et al., 2010). Additionally, the spindle and kinetochore associated Ska complex also recruits PP1 in an RVxF independent manner. All of these PP1-complexes have been suggested to contribute to timely anaphase onset and would be good candidates to promote CDC20 dephosphorylation at attached kinetochores. It should also be considered that at the metaphase-to-anaphase transition most CDC20, together with its kinetochore binding partners BUB1 and BUBR1 (Di Fiore et al., 2015; Lischetti et al., 2014; Vleugel et al., 2015), will have already left the kinetochore. It is therefore conceivable that the bulk of CDC20 dephosphorylation takes place in the cytosol and involves a further, distinct, pool of PP1. The identification of the PP1 pool relevant for CDC20 dephosphorylation is thus an important question to address in the future.

### Timely roles for PP1 and PP2A in APC/C regulation and checkpoint signalling

In different experimental systems PP2A-B55, PP2A-B56 and PP1 have all been reported to act on and regulate components of the APC/C and spindle checkpoint pathways. (Fujimitsu and Yamano, 2020; Hein et al., 2017; Kim et al., 2017). Like PP1, PP2A-B55 is inhibited by CDK1-cyclin B1 and has also been suggested to be an important CDC20 phosphatase in mammalian cells (Hein et al., 2017; Mochida and Hunt, 2012). However, due to the temporal properties of its regulatory mechanism PP2A-B55 only becomes active in anaphase B when cyclin B1 levels fall below a threshold level (Cundell et al., 2013; Cundell et al., 2016). PP1 has been reported to be involved in the activation of PP2A-B55 through the dephosphorylation of the Gwl/MASTL kinase (Heim et al., 2015; Ma et al., 2016; Ren et al., 2017; Rogers et al., 2016), and could thus be considered an upstream regulator of PP2A-B55. PP2A-B55 action on CDC20 is therefore most likely restricted to a later point in anaphase, perhaps at the switch to APC/C regulation by CDH1 and destruction of late anaphase substrates. Consistent with this idea, and in agreement with previous work (Cundell et al., 2013; Hayward et al., 2019a), we did not see any impact on the kinetics of cyclin B1 degradation at the metaphase-to-anaphase transition when PP2A-B55 was depleted (Figure 5).

Unlike PP1 and PP2A-B55, PP2A-B56 is thought to retain full activity in mitosis, and CDK1-dependent phosphorylation of PP2A-B56 recognition sites may in fact increase its activity towards key substrates (Kruse et al., 2013; Smith et al., 2019). Interestingly, a specific pool of PP2A-B56, attached directly to the APC/C, has been implicated in vertebrate CDC20 dephosphorylation during mitosis (Fujimitsu and Yamano, 2020; Lee et al., 2017). Due to the unique properties of PP2A-B56, we propose that this may promote the dynamic turnover of CDC20 phosphorylation during early mitosis, to create a limited level of APC/C activity prior to anaphase (Figure 9). One intriguing possibility is that this is important for the CDC20-dependent but spindle assembly checkpoint independent degradation of cyclin A during mitosis (Di Fiore and Pines, 2010; van Zon and Wolthuis, 2010; Wolthuis et al., 2008). However, at the metaphase-to-anaphase transition, enhanced dephosphorylation of CDC20, promoted by PP1, decisively tips the balance towards APC/C activation, destabilisation of the mitotic state, and consequently mitotic exit. This APC/C^CDC20^ active state is then consolidated by the action of PP2A-B55, the activation of which is initiated by PP1 (Heim et al., 2015; Ma et al., 2016; Ren et al., 2017; Rogers et al., 2016), ensuring that CDC20 is completely dephosphorylated in anaphase (Figure 9).

**Figure 9.**
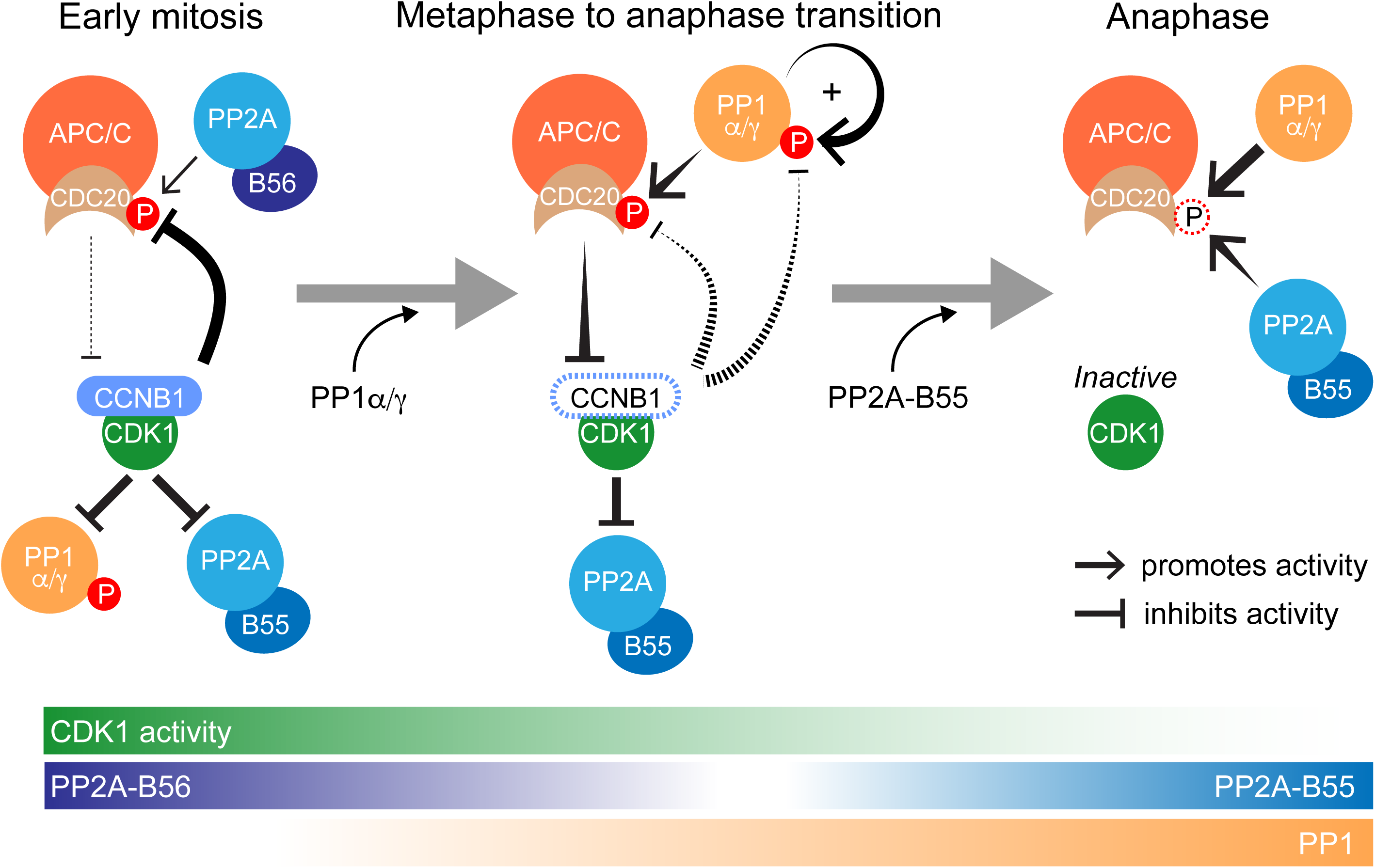
PP2A-B56, PP1 and PP2A-B55 dephosphorylate CDC20 at different points in the cell cycle. Schematic drawing illustrating the dephosphorylation of CDC20 by the three main mitotic phosphatases PP2A-B56, PP1α/γ and PP2A-B55 in early mitosis, at the metaphase-to-anaphase transition and in anaphase, respectively.

We observed that the bulk of spindle checkpoint silencing was carried out with normal kinetics in PP1-depleted cells, or when PP1 was inhibited, presumably by PP2A-B56 (Espert et al., 2014). However, the loss of the last few spindle checkpoint active kinetochores, was significantly delayed under these conditions, confirming a late role for PP1 in the removal of spindle checkpoint proteins from kinetochores (Nijenhuis et al., 2014). Thus, PP1 may be particularly important in promoting cell cycle transitions that have been initiated by PP2A-B56. This provides a compelling reason why PP1 is required at this point in addition to PP2A-B56 to drive the transition into anaphase. The involvement of three major mitotic phosphatases in modulation of CDC20 phosphorylation is thus crucial for restricting APC/C^CDC20^ activity to a narrow window at the metaphase-to-anaphase transition.

## Author contributions

Conceptualisation: F. A. Barr and U. Gruneberg. Investigation: J. Bancroft, J. Holder, Z. Geraghty, T. Alfonso-Pérez, D. Murphy. Funding acquisition: U. Gruneberg, F. A. Barr. Supervision: U. Gruneberg, F. A. Barr. Writing – original draft: U. Gruneberg, F. A. Barr. Writing – review and editing: F. A. Barr and U. Gruneberg with input from all authors.

## Acknowledgements

JB and ZG were supported by a Medical Research Council Senior Non-Clinical Research fellowship awarded to UG (MR/K006703/1), and TAP by a Biotechnology and Biological Sciences Research Council Strategic LoLa grant (BB/M00354X/1). JH was supported by a Wellcome Trust PhD award and a Cancer Research UK program grant award (C20079/A15940) to FAB. We acknowledge Catherine Wormald and Flavia Scialpi for initial observations, Fengying Liu for help with the generation of CDC20 Flp-In TRex cell lines and Renaud Caous and Emile Roberts for help with live cell imaging analysis. We thank Elena Poser and Dan Hayward for critical reading of the manuscript.

## Material and Methods

### Chemicals and antibodies

General laboratory chemicals and reagents were obtained from Sigma-Aldrich and Thermo-Fisher Scientific. Drugs were dissolved in DMSO unless specifically indicated. Inhibitors were obtained from Sigma Aldrich (CDK1 inhibitor flavopiridol, 5 mM stock), Tocris Bioscience (MPS1-inhibitor AZ3146 20 mM stock; PP1 and PP2A-inhibitor calyculin A 1 mM stock, PP1-inhibitor tautomycetin, 2.5 mM stock), Insight Bioscience (proteasome inhibitor MG132 20 mM stock) and Merck (microtubule polymerisation inhibitor nocodazole 0.66 mM stock). Thymidine (Sigma Aldrich, 100mM stock) and doxycycline (Invivogen, 2 mM stock) were dissolved in water. Commercially available polyclonal (pAb) or monoclonal (mAb) antibodies were used for beta Actin (HRP conjugated; Mouse mAb Abcam, [AC-15] ab49900), Tubulin (Mouse mAb; Sigma, [DM1A] T6199), PPP1CA (Rabbit pAb; Bethyl, A300-904A), PPP1CA-pT320 (Rabbit pAb, Abcam, Ab62334), PPP1CB (Rabbit pAb, Bethyl, A300-905A), PPP1CC (Goat pAb; Santa Cruz, sc6108), Securin (Rabbit pAb, Abcam, Ab79546), CDC27/APC3 (mouse mAb clome C-4, Santa Cruz sc13154), CDC20 (mouse mAb clone E-7, Santa Cruz sc-13162; mouse mAB, clone AR12, Millipore UK Ltd, MAB3775; mouse mAb, clone BA8, Bio-Techne (R&D systems), NB100-2646 – these three antibodies were mixed for immunoprecipitations. CDC20 (rabbit pAb, Santa Cruz, sc-8358) was used for Western blotting.

Antibodies against CDC20-pT70 were raised in rabbits using phospho-peptide CSKVQT(pT)PSKPG and affinity-purified using an immobilised form of the same peptide (Moravian). Secondary donkey antibodies against mouse, rabbit, guinea pig or sheep and labelled with Alexa Fluor 488, Alexa Fluor 555, Alexa Fluor 647, Cy5, or HRP were purchased from Molecular Probes and Jackson ImmunoResearch Laboratories, Inc., respectively. Affinity purified primary and HRP-coupled secondary antibodies were used at 1 µg/ml final concentration. For western blotting, proteins were separated by SDS-PAGE and transferred to nitrocellulose using a Trans-blot Turbo system (Bio-Rad). Protein concentrations were measured by Bradford assay using Protein Assay Dye Reagent Concentrate (Bio-Rad). All western blots were revealed using ECL (GE Healthcare).

### Molecular biology and siRNA reagents

Human CDC20 was amplified from human testis cDNA (Marathon cDNA; Takara Bio Inc.) using Pfu polymerase (Agilent Technologies). CDC20 expression constructs were made using pcDNA5/FRT/TO vectors (Invitrogen) modified to encode the EGFP or FLAG reading frames. Mutagenesis to introduce phospho-site mutations and resistance to CDC20 siRNA oligo #14 was performed using the QuikChange method (Agilent Technologies). DNA primers were obtained from Invitrogen. For the knock down of PPP1CA and PPP1CC small interfering RNA (siRNA) duplexes 5’-UGGAUUGAUUGUACAGAAAUU-3’ and 5’-GCGGUGAAGUUGAGGCUUAUU-3’ targeting the 3’-UTR of PPP1CA and PPP1CC, respectively were used in Figures 2, 3 and 6 and 5’-CAUCUAUGGUUUCUACGAU-3’ and 5’-GAACGACCGUGGCGUCUCU-3’ (for PPP1CA); and 5’-GCGGAGAGUUUGACAAUGC-3’ and 5’-UAGAUAAACUCAACAUCGA-3’ (for PPP1CC), targeting the ORFs, were used in Figures 5, 7 and Figure S2. PPP1CB was depleted using a 3’-UTR oligo 5’-GGGAAGAGCUUUACAGACAUU-3’. CDC20 was depleted using siRNA oligo #14 5’-CGGAAGACCUGCCGUUACA-3’ (ThermoFisher). siRNA oligos for PP2A-B55 and PP2A-B56 have been described (Hayward et al., 2019a; Hayward et al., 2019c).

### Cell culture and CRISPR procedures

HeLa cells were cultured in DMEM with 1% [vol/vol] GlutaMAX (Life Technologies) containing 10% [vol/vol] bovine calf serum at 37°C and 5% CO_2_. For plasmid transfection and siRNA transfection, Mirus LT1 (Mirus Bio LLC) and Oligofectamine (Invitrogen), respectively, were used. HeLa cell lines with single integrated copies of the desired transgenes were created using the T-Rex doxycycline-inducible Flp-In system (Invitrogen)(Tighe et al., 2004)). CRISPR/Cas9-edited HeLa cells with an inserted GFP tag in the C-terminus of the CCNB1 gene product and HeLa cells stably expressing GFP-MAD2 have been described before (Alfonso-Perez et al., 2019; Hayward et al., 2019a). To enable visualisation of CCNB1 in the HeLa Flp-In TRex background, CCNB1-mCherry was integrated into the endogenous CCNB1 locus of parental HeLa Flp-In TRex cells as described (Alfonso-Perez et al., 2019).

### High-resolution mitotic exit time courses for Western blotting

HeLa cells were seeded at 1,200,000 cells/dish onto 15 × 15 cm dishes per condition and grown for 72 hr. Nocodazole, a microtubule depolymerisation agent, was added to 100 ng/ml (330 nM), for 20 hr, to arrest the cells in mitosis. If depletion was necessary, cells were seeded at 600,000 cells/dish and grown for 24 hr before transfection with the appropriate siRNA for 72 hr. Nocodazole was added to the dishes for the final 20 hr of depletion. The mitotic cells were harvested by shake off, washed in 2x 25 ml 1x PBS, 1x 25 ml Opti-MEM, both of which had been pre-equilibrated to 37°C, 5% CO_2_. Wash centrifugations were carried out for 5 min at 200 g_av_, 37°C. The cells were then resuspended in pre-equilibrated Opti-MEM to give 15,000,000 cells/ml. The cells were incubated for 25 min (37°C, 5% CO_2_) to allow cells to rebuild bipolar mitotic spindles, with gentle mixing every ∼5 min. While the cells were incubating, 500 μl of flavopiridol buffer (480 µl pre-equilibrated Opti-MEM + 20 µl, 5 mM flavopiridol) was made and pre-warmed to 37°C. Where more than 500 µl of flavopiridol was required, this was scaled accordingly. Flavopiridol buffer was added to the cells at a 1:10 dilution, giving a final concentration of 20 µM. The cells were immediately mixed through inversion and pipetting before being split in half. Owing to the frequency of early timepoints, one 2 ml aliquot was kept incubating at 37°C, 5% CO_2_, gently mixed every 5 min, to ensure stable conditions. The other half was placed in a 37°C water bath and sampled accordingly. At each timepoint, 50 µl of cells was added to 25 µl of 3x sample buffer and boiled for 5 minutes. Samples were diluted 1:2 with 1x sample buffer prior to Western blotting. Typically, 6 µg was loaded on each Western blot but when necessary was increased to 10 µg as required. Where necessary, drugs were added at the start of the 25 min incubation, to ensure maximal inhibition prior to the addition of flavopiridol. For MPS1 inhibition time courses (and for the comparative CDK1 inhibition time course in Figure S1), drug treatments and washes for nocodazole release were performed in complete media (DMEM with 1% [vol/vol] GlutaMAX (Life Technologies) containing 10% [vol/vol] bovine calf serum) at 37°C and 5% CO_2_. MPS1 inhibitor (MPS1-i; AZ3146) was pre-diluted in complete media and pre-warmed to 37°C before being added to cell suspensions (1.5×10^7^ cells/ml) to a final concentration of 2 µM. For sample collection, cells were treated with 50 nM calyculin A to prevent further de-phosphorylation events during centrifugation and 1x PBS wash, before being re-suspended in lysis buffer supplemented with phosphatase and protease inhibitors for snap-freezing.

### Analysis of CDC20 phosphorylation

For analysis of CDC20 phosphorylation 10% [wt/vol] polyacrylamide separating gels were prepared with 100 µM MnCl2 and 25 µM PhosTag reagent (Wako Chemicals, AAL-107S1). Double concentrations of APS and TEMED were used to aid polymerisation. Typically, 10 µg of lysate was loaded for PhosTag gels. Prior to transfer, gels were equilibrated with 3x 3 min washes in transfer buffer (20 mM Tris; 150 mM Glycine; 0.1% [wt/vol] SDS; 20% [vol/vol] MeOH; 20 mM EDTA) and subsequently 3x 3 min washes in transfer buffer without EDTA (20 mM Tris; 150 mM Glycine; 0.1% [wt/vol] SDS; 20% [vol/vol] MeOH).

Analysis with anti-CDC20-pT70 antibodies was carried out on CDC20 immunoprecipitates. For CDC20 immunoprecipitations, 1.8×10^6^ cells per sample were lysed for 15 min on ice, vortexing briefly every 5 min, in lysis buffer supplemented with phosphatase and protease inhibitors (20 mM Tris-HCl pH 7.5; 150 mM NaCl; 1 % IGEPAL [vol/vol]; Phosphatase inhibitor cocktail 1:100 [Sigma-Aldrich]; Protease inhibitor cocktail 1:250 [Sigma-Aldrich]; 0.5 M β-glycerol phosphate; 10 mM NaF; 100 nM Okadaic acid; 100 nM Calyculin A; PMSF 1 mM). Volumes of lysis buffer were calculated during sample collection to give lysate concentrations of ∼1 mg/ml. Cells were then centrifuged at 14,000 g_av_ for 15 minutes at 4°C to produce a cleared lysate which was analysed by Bradford assay. For the experiment in Figure 6C, CDC20 was isolated from 0.35 mg of lysate by 90 min incubation at 4°C with 20 µl Protein-G Dynabeads (ThermoFisher, 10004D) and 2 µg of each of three anti-CDC20 mAbs: clone E-7, clone BA8, clone AR-12. For the experiment in Figure 6G, CDC20 was isolated from 1 mg of lysate by 1 h incubation at 4°C with 40 µl Protein-G Dynabeads and 3 µg mAb clone E-7. Dynabeads were washed 3x with lysis buffer and 3x with wash buffer (20 mM Tris-HCl pH 7.5; 150 mM NaCl; 0.1 % IGEPAL [vol/vol]) and re-suspended in 100 µl 2.5x Laemmli sample buffer. For immunoprecipitation of GFP-CDC20, 20 µl Protein-A Dynabeads were used with 2.5 µg anti-GFP (rb pAb, Abcam ab290), or anti-mCh (rb pAb, Abcam ab167453) for control.

### Functional analysis of CDC20 and CDC20^6A^

For CDC20 siRNA rescue experiments, HeLa Flp-In TRex cells expressing CCNB1-mCherry from the endogenous promoter and GFP-CDC20^WT^ or GFP-CDC20^6A^ from the Flp-In site were used. CDC20 siRNA rescue was performed by induction with 2 µM doxycycline of GFP-CDC20 transgenes (WT and 6A) resistant to siRNA oligo #14, for 6 hours prior to a 48-hour siRNA depletion of endogenous CDC20 using oligo #14. A second induction was performed 18 hours into the siRNA depletion. For live cell imaging and sample collection for immunoprecipitation, cells were treated with 2 mM thymidine 18 hrs after RNAi addition, for 18 hours. The thymidine was removed by washing 3 times with DMEM, with 2 µM doxycycline re-added in the final wash. For immunoprecipitation samples, 330 nM nocodazole was added 8 hours after thymidine release for 4 hours, and cells were collected by mitotic shake-off. For live cell imaging SiR-DNA (Spirochrome) was added to the final wash of the thymidine release at a concentration of 100nM, and imaging commenced 9 hours later.

### Live cell imaging of MAD2 and CCNB1

For the live cell imaging of GFP-MAD2 CCNB1-mCherry cells, cells were seeded into 35-mm dishes with 1.5-thickness coverglass bottom (Fluorodishes - Applied Precision) at ∼50,000 cells per dish and grown for 24 hours. Cells were then RNAi depleted for a total of 60 hours. 35 hrs after RNAi addition, cells were treated with 2 mM thymidine for 18 hours. The thymidine was removed by washing 3 times with 2 ml of DMEM and imaging was started 9 hours later. SiR-DNA (Spirochrome) was added to the final wash at a concentration of 100nM. In assays using PP1 inhibitor, 5 µM tautomycetin was added to cells 30 min before imaging started. Live cell imaging was performed using a spinning disc confocal system (Ultraview Vox; PerkinElmer) mounted on an inverted microscope (IX81; Olympus) equipped with an EM charge-coupled device camera (C9100-13; Hamamatsu Photonics) and controlled by Volocity software, except for CDC20 RNAi rescue assays which were imaged on a Deltavision Elite system using an inverted microscope (IX81; Olympus) and equipped with a QuantEM EMCCD camera (Photometrics). Cell were placed in a 37°C and 5% CO2 environmental chamber (Tokai Hit) on the microscope stage with lens heating collar. Imaging was performed using a 60× NA 1.4 oil immersion objective lens.

To monitor chromosome congression, checkpoint silencing and CCNB1 degradation, HeLa CCNB1-mCherry GFP-MAD2 cells treated with SiR-DNA at 100 µM were imaged using 8% 561 nm laser power with 100 ms exposure for CCNB1-mCherry, 6% 488 nm laser power with 80 ms exposure for GFP-MAD2 and and 2% 647 nm laser power with 20ms exposure for SiR-DNA. 19 axial planes were captured at 0.6 μm apart (Z then wavelength) at an interval of 2 min for 10h. These images were then used to determine last chromosome congression time and the time at which the last MAD2 foci disappeared. The levels of CCNB1 were observed relative to the checkpoint status.

### Spindle checkpoint silencing in live cells

For live cell imaging with the addition of the MPS1 inhibitor AZ3146, HeLa cells expressing GFP-MAD2 and CCNB1-mCherry were treated as above but seeded into imaging dishes covered with lids containing a preformed hole to facilitate drug addition. Cells were imaged at intervals of 2 min and cells in mitosis were visually scanned to identify cells approaching metaphase. After 4 captured time points, MPS1-i diluted in 200 µl DMEM was added to the cells to a final concentration of 2 µM. After drug addition, imaging was continued for 1-2hr.

### Statistical analysis

Statistical analysis of live cell imaging data and intensity measurements was carried out in Excel and GraphPad Prism. Statistical tests in Figure 3 were unpaired T-test with Welch’s correction for unequal SD and Mann-Whitney tests in Figure 8. Unless stated otherwise, the quantitations are derived from a compilation of three independent experiments. For Western blot analysis, representative examples of three independent repeats are shown.

