## Supplemental figures for "PP1 promotes cyclin B destruction and the metaphase-anaphase transition by dephosphorylating CDC20"

#### Supplemental Figure Legends

**Figure S1.** CDK1 inhibition accelerates cyclin B1 degradation at the metaphase-to-anaphase transition. **(A)** Synchronous progression of mitotic HeLa cells into anaphase was triggered with CDK1 inhibitor (5 µM flavopiridol) or **(B)** MPS1 inhibitor (2 µM AZ3146), and samples were collected every 5 min and blotted with the indicated antibodies. **(C)** Densitometric measurement and line graph of cyclin B1 levels in (A) and (B). **(D)** Densitometric measurement and line graph of PP1 pT320 levels in (A) and (B). **(E)** HeLa-Flp-In TRex cells induced to express PP1α-GFP or PP1γ-GFP were depleted of PP1α/γ and processed for immunofluorescence analysis. CENP-A and KNL1 were stained as kinetochore references. **(F)** PP1α-GFP or PP1γ-GFP kinetochore intensity in control or PP1α/γ depleted cells was quantified relative to CENP-A. **(G)** Western blot analysis of PP1α/γ depleted cells. Tubulin was used as a loading control.

**Figure S2.** PP1 regulates cyclin B destruction downstream of checkpoint silencing. **(A)** HeLa cells stably expressing GFP-MAD2 were treated with control, **(B)** siPP1α/γ or **(D)** siPP2A-B56 for 60 hours, or **(C)** 5 µM tautomycetin PP1-inhibitor for 30 min before imaging. DNA was visualised with SiR-Hoechst DNA dye. Cells were imaged every 2 mins for a total of 10 hrs. **(E)** Total cellular intensity of kinetochore-localised MAD2 ($MAD2^{KT}$) is plotted as a function of time for siControl, siPP1α/γ and PP1-i (n=6), and siPP2A-B56 (n=5). **(F)** Bar graphs showing the mean time interval ± SD between the last chromosome congressed (LCC) and last MAD2 lost (LML), and last MAD2 lost and anaphase onset (ANA), in control (n=60) and siPP1α/γ cells (n=34) expressing GFP-MAD2.

**Figure S3.** Anti-CDC20-pT70 antibodies specifically recognise phosphorylated CDC20. **(A)** HeLa Flp-In TRex cells depleted of the endogenous CDC20 and expressing GFP-CDC20$^{WT}$ or GFP-CDC20$^{6A}$ were treated with nocodazole for 20 min, then stained for CDC20 pT70. **(B)** CDC20 pT70 normalised for total GFP-CDC20 is plotted as mean ± SD. **(C)** HeLa Flp-In TRex cells depleted of the endogenous CDC20 were induced for GFP-CDC20$^{WT}$ or GFP-CDC20$^{6A}$, then arrested in mitosis. GFP-CDC20 was immunoprecipitated using anti-GFP antibodies (IP), and the immunoprecipitated samples Western blotted for total CDC20, CDC20 pT70 and APC3. APC3 precipitates more strongly with non-phosphorylatable CDC20 (GFP-CDC20$^{6A}$) than with mitotically phosphorylated CDC20, as previously reported (Hein and Nilsson, 2016). **(D)** Western blots confirming depletion of CDC20 and PP2A-B55.

### Figure S1

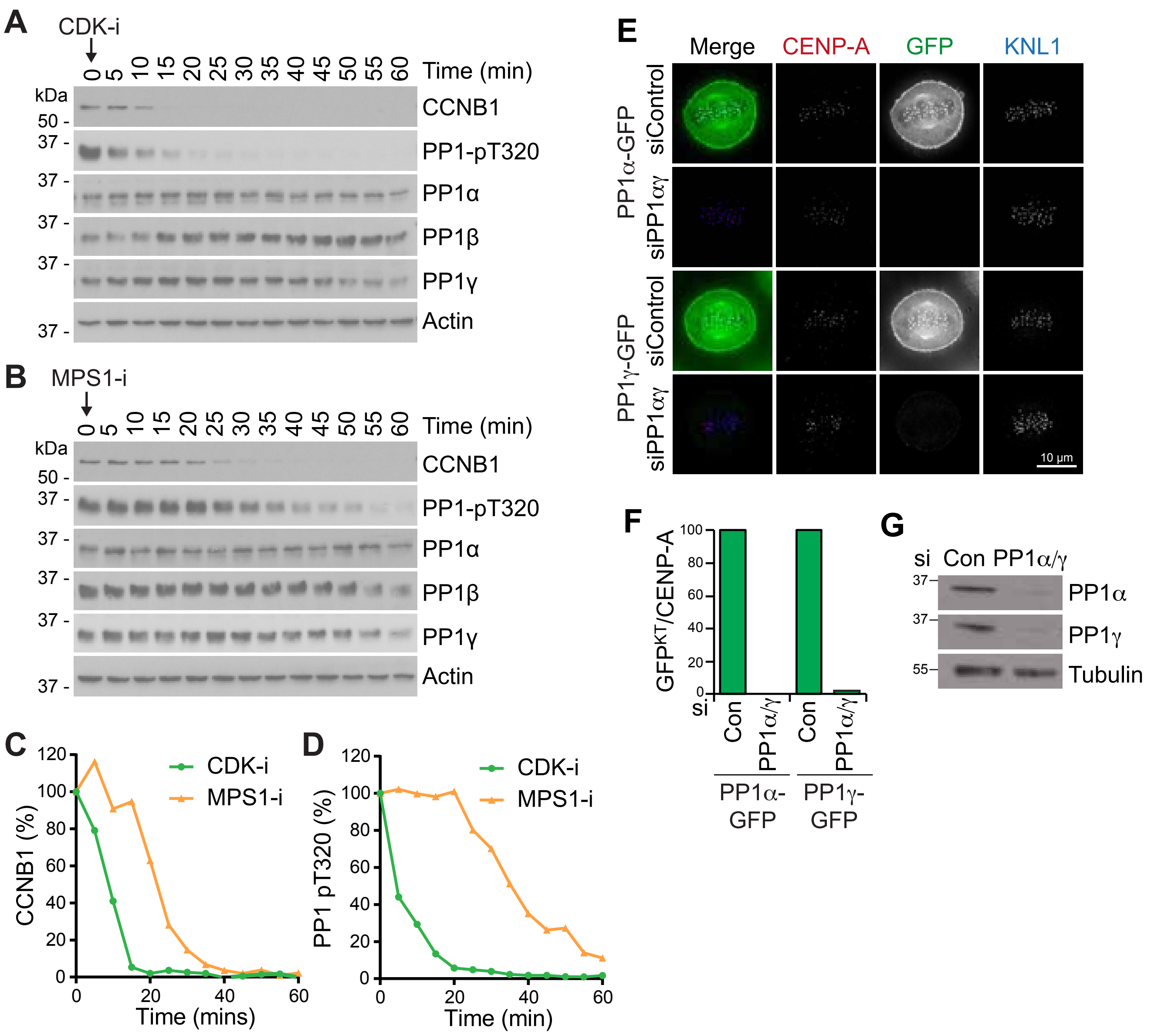
A
CDK-i
Time (min)
0 5 10 15 20 25 30 35 40 45 50 55 60
kDa
50 -
37 -
CCNB1
PP1-pT320
PP1α
PP1β
PP1γ
Actin
B
MPS1-i
C
CCNB1 (%)
Time (mins)
CDK-i
MPS1-i
D
PP1 pT320 (%)
Time (min)
E
Merge
CENP-A
GFP
KNL1
PP1α-GFP
PP1γ-GFP
siControl
siPP1αγ
10 µm
F
GFP^KT/CENP-A
si
Con
PP1α/γ
PP1α-GFP
PP1γ-GFP
G
si Con PP1α/γ
PP1α
PP1γ
Tubulin

### Figure S2

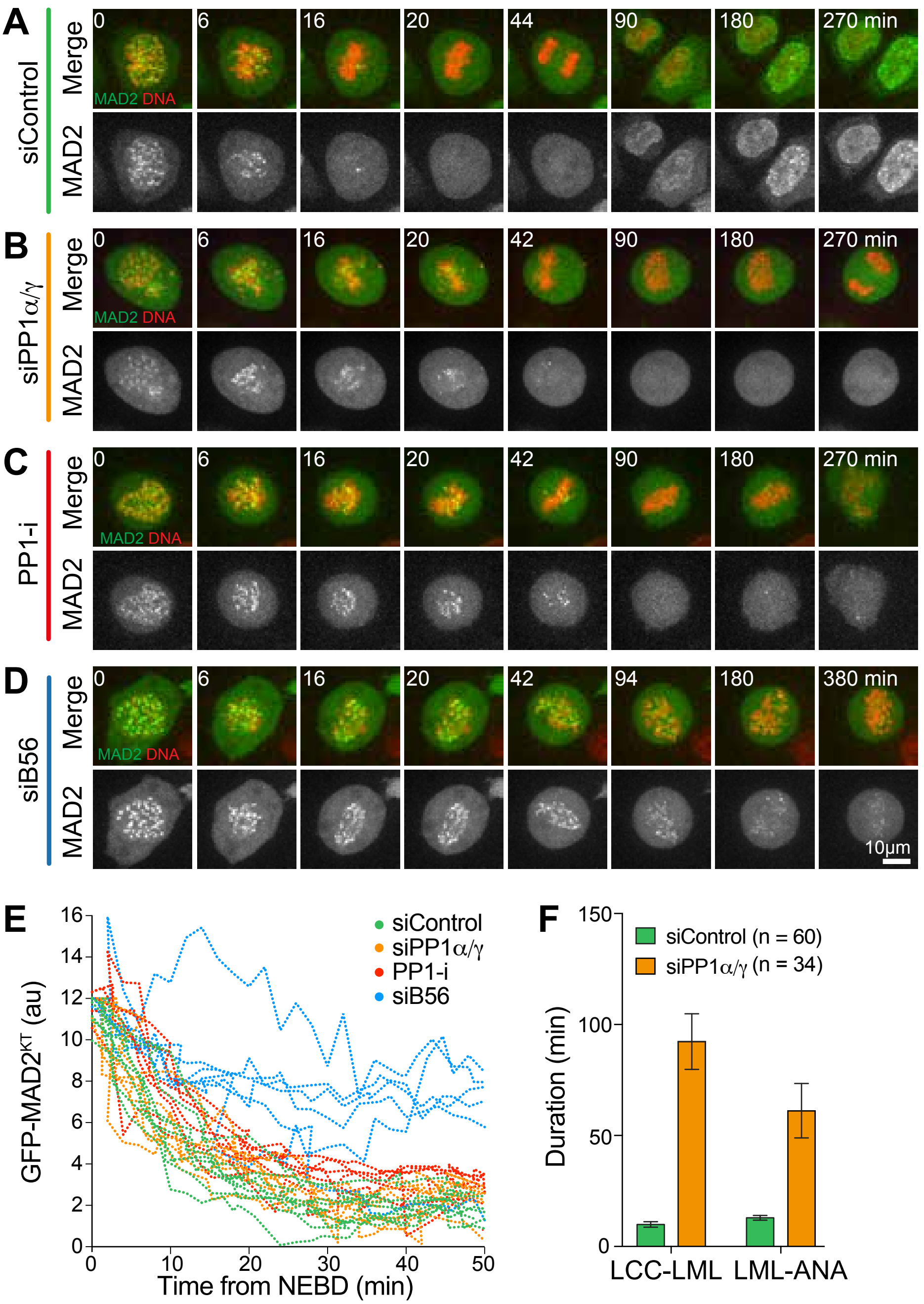
A
siControl
Merge
MAD2
0
6
16
20
44
90
180
270 min
MAD2 DNA
B
siPP1α/γ
Merge
MAD2
0
6
16
20
42
90
180
270 min
MAD2 DNA
C
PP1-i
Merge
MAD2
0
6
16
20
42
90
180
270 min
MAD2 DNA
D
siB56
Merge
MAD2
0
6
16
20
42
94
180
380 min
MAD2 DNA
10μm
E
GFP-MAD2KT (au)
Time from NEBD (min)
siControl
siPP1α/γ
PP1-i
siB56
0
2
4
6
8
10
12
14
16
0
10
20
30
40
50
F
Duration (min)
siControl (n = 60)
siPP1α/γ (n = 34)
0
50
100
150
LCC-LML
LML-ANA

#### Figure S3

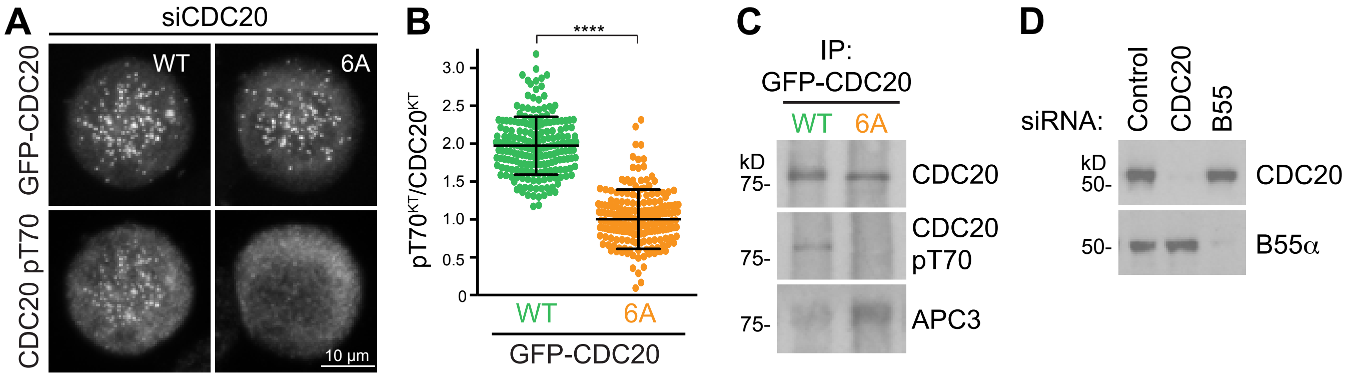
A
siCDC20
GFP-CDC20
CDC20 pT70
WT
6A
10 µm
B
****
pT70KT/CDC20KT
3.0
2.5
2.0
1.5
1.0
0.5
0
WT
6A
GFP-CDC20
C
IP:
GFP-CDC20
WT
6A
kD
75-
75-
75-
CDC20
CDC20
pT70
APC3
D
siRNA:
Control
CDC20
B55
kD
50-
50-
CDC20
B55α
